# CRISPR-Cas is associated with fewer antibiotic resistance genes in bacterial pathogens

**DOI:** 10.1101/2021.04.12.439454

**Authors:** Elizabeth Pursey, Tatiana Dimitriu, Fernanda L. Paganelli, Edze R. Westra, Stineke van Houte

**Affiliations:** Environment and Sustainability Institute, Biosciences, University of Exeter, Penryn, Cornwall, United Kingdom; Department of Medical Microbiology, University Medical Center Utrecht, Utrecht, The Netherlands

**Keywords:** CRISPR-Cas, horizontal gene transfer, mobile genetic elements, antibiotic resistance, plasmids, integrative conjugative elements

## Abstract

The acquisition of antibiotic resistance genes via horizontal gene transfer is a key driver of the rise in multidrug resistance amongst bacterial pathogens. Bacterial defence systems per definition restrict the influx of foreign genetic material, and may therefore limit the acquisition of antibiotic resistance. CRISPR-Cas adaptive immune systems are one of the most prevalent defences in bacteria, found in roughly half of bacterial genomes, but it has remained unclear if and how much they contribute to restricting the spread of antibiotic resistance. We analysed ~40,000 whole genomes comprising the full RefSeq dataset for 11 species of clinically important genera of human pathogens including *Enterococcus*, *Staphylococcus*, *Acinetobacter* and *Pseudomonas*. We modelled the association between CRISPR-Cas and indicators of horizontal gene transfer, and found that pathogens with a CRISPR-Cas system were less likely to carry antibiotic resistance genes than those lacking this defence system. Analysis of the mobile genetic elements targeted by CRISPR-Cas supports a model where this host defence system blocks important vectors of antibiotic resistance. These results suggest a potential “immunocompromised” state for multidrug-resistant strains that may be exploited in tailored interventions that rely on mobile genetic elements, such as phage or phagemids, to treat infections caused by bacterial pathogens.

## Introduction

The spread of antibiotic resistance (ABR) genes between bacterial strains and species is of huge global importance; without access to working antibiotics, much of modern medicine is threatened, including surgery, cancer treatment and neonatal care [1]. Opportunistic pathogens in particular often have the ability to take up DNA readily from the environment, through various mechanisms of horizontal gene transfer (HGT) [2]. In some cases, strains are naturally competent and therefore able to uptake extracellular DNA from their environment, as occurs in some clinical isolates of *Acinetobacter baumannii* [3]. Mobile genetic elements (MGEs) also enter the cell through transduction (transfer of DNA by bacteriophages), or conjugation, in which mobile elements are physically transferred between cells through direct cell-cell contact. Conjugative plasmids encode all necessary self-transfer machinery, while mobilizable plasmids require it to be encoded by another element, but both contribute hugely to the spread of ABR genes. For example, conjugative plasmids are enriched for ABR genes and are able to transfer these resistance determinants across species and genera [4]. In addition, they often harbour integrons, characterised by the *intI1* integrase gene, which enables them to capture ABR and other genes in cassettes that can be spread by the conjugative plasmid [5]. Integrative conjugative elements (ICEs) are widespread in bacteria, and more numerous than conjugative plasmids [6]. These MGEs can integrate into the host chromosome and transmit vertically through cell division, but can also excise themselves and be transferred horizontally through conjugation [7].

Host defences may block the acquisition of MGEs. One of the most prevalent defences is CRISPR-Cas (<u>c</u>lustered <u>r</u>egularly <u>i</u>nterspaced <u>s</u>hort <u>p</u>alindromic <u>r</u>epeat (CRISPR) loci and <u>C</u>RISPR-<u>as</u>sociated (*cas*) genes), an adaptive immune system found in in ~30-40% of bacteria [8,9], in which short spacer sequences, derived from incoming DNA, are incorporated at CRISPR loci in between repeat sequences. Functioning as a form of immunological “memory”, these spacer sequences guide their cognate Cas enzymes to cut and destroy any sequence matching this immune record. This enables the cell to defend itself from predation by bacteriophages, and block other sources of potentially costly incoming MGEs.

A number of studies have examined the extent to which CRISPR-Cas restricts HGT, using genomic approaches. In contrast to virus-mediated immunity, which is often circumvented through viral mutation, overcoming immunity to elements such as plasmids often involves loss or inactivation of the CRISPR-Cas system itself [10]. This means evidence for its potential role in blocking these elements may be found in the genomic and phylogenetic distributions of CRISPR-Cas systems in bacteria.

For example, some studies have examined whether CRISPR-Cas systems prevent HGT by looking at genome length. Theoretically, genomes possessing CRISPR-Cas may be shorter if they are blocking acquisition of foreign DNA elements. Evidence for this has been found in *Pseudomonas aeruginosa* [11,12], a species with a large core and accessory genome. More direct interactions between HGT and CRISPR-Cas can be tested using genomic comparisons. However, the outcomes of these studies have been ambiguous. For example, a study of 1399 bacterial and archaeal genomes did not reveal a correlation between markers of recent HGT and spacer count (used as a proxy for CRISPR-Cas activity) [13]. However, a more recent analysis compared pairs of conspecific genomes with and without CRISPR-Cas as a method to control for relatedness, and found fewer plasmids when CRISPR-Cas was present within 29 genome pairs [14]. When comparing over 100,000 genomes from multiple species, another study revealed a complex situation when looking for correlations between particular ABR gene classes and CRISPR-Cas presence, in which positive and negative relationships were found dependent on species and ABR gene [15].

Research focused on within-species comparisons has often found more compelling evidence for CRISPR-Cas as a barrier to HGT and the spread of ABR genes. In *P. aeruginosa*, in paired within-ST (sequence type) genomes, fewer ICEs and prophages were detected when CRISPR-Cas was present [12], whilst another study found negative associations between the presence of type I-E and I-F systems and particular ABR genes [11]. Studies in Enterococci also support the role of CRISPR-Cas in blocking ABR acquisition, with fewer resistance genes detected in genomes with CRISPR-Cas in a set of 48 strains of *E. faecalis* and 8 strains of *E. faecium* [16]. Further genomic associations between CRISPR-Cas presence and a lack of ABR genes has been detected in *Klebsiella pneumoniae* [17,18], whilst there is evidence in *A. baumannii* that CRISPR-Cas presence is correlated with an absence of plasmids [19].

Here, we sought to expand on existing work using the complete RefSeq dataset for a group of bacterial human pathogens from 11 species (n=39,511), with diverse lifestyles, and CRISPR-Cas system types. We built upon the null hypothesis significance testing used in previous studies, which typically used *t*-tests (or their non-parametric alternatives) or correlation analyses. To do so, we applied linear and generalised linear models (LMs, GLMs) with model selection based on Akaike information criterion (AIC) values, which allows comparison of all candidate models and selection of the model with the best fit to the dataset. We also used Bayesian phylogenetic models to examine within-species relationships, to control for genetic distance between genomes. Both of these model types are advantageous in that they allow predictions to be made about the dataset based on values of different variables.

We tested both between and within-species associations between ABR gene counts and CRISPR-Cas presence. First, we assessed whether the probability that a genome possesses a CRISPR-Cas system changes based on the presence of plasmids, ICEs and integrons, as well as looking at whether genomes with CRISPR-Cas were shorter. Subsequently, we built within-species models, controlling for genetic distance, to detect associations between specific CRISPR-Cas types and spacer counts, and the quantity of ABR genes accumulated in a genome. Finally, we assessed whether a CRISPR-Cas system possessing spacers that target MGEs that are vectors of ABR genes can effectively reduce the ABR count in the genome.

## Methods

### Genomes

RefSeq genomes for *P. aeruginosa*, *A. baumannii*, *Neisseria meningitidis*, *Staphylococcus epidermidis*, *Streptococcus pyogenes*, *Francisella tularensis*, *Mycobacterium tuberculosis*, *Neisseria gonorrhoeae*, *E. faecium* and *E. faecalis* were retrieved from NCBI between December 2020 and January 2021 in nucleotide fasta format using ncbi-genome-download v0.3.0 (https://github.com/kblin/ncbi-genome-download). The strains were selected to include a variety of CRISPR-Cas system types, pathogenic lifestyles (i.e. obligate pathogens such as *F. tularensis*, *M. tuberculosis* and *N. gonorrhoeae* as well as opportunistic pathogens like *P. aeruginosa*, *S. aureus and S. pyogenes*), and species of significance in the spread of ABR such as *A. baumannii* and *E. faecium*. The species used are summarised in Table 1.

**Table 1 –.** Summary of species included

| Species | Abbreviation used | RefSeq genomes (n) |
| --- | --- | --- |
| *Pseudomonas aeruginosa* | PA | 5360 |
| *Acinetobacter baumannii* | AB | 4864 |
| *Neisseria meningitidis* | NM | 1967 |
| *Staphylococcus epidermidis* | SE | 908 |
| *Staphylococcus aureus* | SA | 12148 |
| *Streptococcus pyogenes* | SP | 2106 |
| *Francisella tularensis* | FT | 675 |
| *Mycobacterium tuberculosis* | MT | 6688 |
| *Neisseria gonorrhoeae* | NG | 723 |
| *Enterococcus faecium* | EFm | 2247 |
| *Enterococcus faecalis* | EFs | 1825 |
| Total |  | 39511 |

### *Identification of CRISPR-Cas systems, ABR genes, plasmids, ICEs, and* intI1

CRISPR-Cas systems were detected using CRISPRCasTyper [20], and orphan CRISPR arrays were not included in any spacer analysis. CRISPR-Cas systems with “Unknown” predictions were also removed from the dataset. Acquired ABR genes and plasmid replicons were identified using abricate (https://github.com/tseemann/abricate) with the NCBI AMRFinderPlus [21] and PlasmidFinder [22] databases, both from 19/04/2019 and containing 5386 and 460 sequences, respectively. To detect ICEs and *intI1*, custom BLAST databases were created based on the ICEberg database (retrieved January 2021; Liu et al. 2019), and the NCBI gene entry for class 1 integron integrase *intI1* [NC_019069.1 (99852.100865, complement)], and used with abricate.

### Identification of spacer targets and MGEs carrying ABR genes

Spacer sequences were searched against BLAST databases constructed from the aforementioned ABR, ICE and prophage databases as well as a curated plasmid database [24], and the complete RefSeq viral release (retrieved February 2021). Only hits with 100% identity were used for downstream analyses. The total number of spacers per genome was approximated by subtracting 1 from the number of repeats for each array and summing these values, and any spacers with no hits in any database were assigned as “unknown”.

MGEs carrying ABR genes were identified by running a blastn search with default parameters, querying all sequences in search databases used for ICE, plasmid and virus detection against a custom BLAST database built from the aforementioned NCBI AMRFinderPlus database. Subsequently, a percent identity cutoff of 90% was applied to hits. The presence of ABR genes on MGEs was classified in a binomial way, in which ≥ 1 ABR gene was counted as the presence of ABR on the element. In cases where multiple MGEs were targeted by a single spacer, the spacer was classified as targeting an MGE with ABR if at least one of these MGEs carried at least one ABR gene.

### Data analysis and statistical modelling

The University of Exeter’s Advanced Research Computing facilities were used to carry out this work. Data analyses were conducted in R v4.0.2 using the tidyverse suite of packages [25]. Unless otherwise specified, linear or generalized linear models were fitted using the *lm* or *glm* functions in base R. Linear models allow the inclusion of multiple predictors in one model, termed fixed effects, to estimate the relationship between these predictors and a response variable. These models generate a formula that calculates the mean of a response variable based on the value of each fixed effect (predictor). Regression coefficients are fitted, which give the slope and the intercept (value of the response variable when the predictor = 0) for each fixed effect. For categorical predictors, a different intercept is fitted for each level of the predictor (e.g. each species in a multispecies model). To fit different slopes for levels of categorical predictor, interaction terms must be fitted between them and a continuous predictor. For example, an interaction between species and the count of ABR genes would allow varying slopes and intercepts for each individual species.

For these analyses, a maximal model was generated and all possible candidate models were compared using the AIC method using *dredge* from the MuMIn package [26]. AIC values assess the fit of a model by looking at the likelihood of a model given the data, penalising for increased number of parameters (as increased complexity of the model increases parameter uncertainty). The model with the lowest AIC value was selected, and no alternative models were within 2 AICs of the best fitting model.

Prediction data frames with 95% confidence intervals were generated using *ggpredict* from the package ggeffects [27], model dispersion was tested and scaled residuals were examined using DHARMa residual diagnostics [28], and the final predictions were visualised with ggplot2, ghibli [29] and cowplot (https://wilkelab.org/cowplot/index.html). Prediction plots were produced only for biologically realistic combinations of fixed effects and response, i.e. those where sufficient data was present in the data set.

Associations between genome length and CRISPR-Cas presence/absence were tested using a linear model with genome length as the response variable and CRISPR-Cas presence/absence and species as fixed effects, with CRISPR-Cas presence/absence × species as an interaction term.

The effect of various MGEs on the distribution of CRISPR-Cas was modelled using a binomial GLM with CRISPR-Cas presence/absence as the response variable and counts of ICEs, plasmid replicons, ABR genes and *intI1* copies, as well as species, as fixed effects. In addition, we fitted an interaction between species and every other fixed effect. Prediction data frames were created up to the maximum number of each MGE detected in each species (unless this value was an outlier, in which case a more biologically informative upper limit was set).

The relationship between spacers targeting MGEs with and without ABR and the count of ABR genes was modelled using a Bayesian Poisson GLM in MCMCglmm. The default weakly informative priors were used, and models were run for 20,000 iterations with a burn in period of 10,000 iterations and a thinning interval of 5. Two models were run, with the number of ABR genes as the response variable, species as a fixed effect, and either the counts of spacers targeting MGEs with ABR or targeting MGEs without ABR as an additional fixed effect. Prediction data frames were created up to the maximum number of spacers detected targeting MGEs with and without ABR.

### Construction of trees and genetic distance-controlled single-species models

The package mashtree [30] with mash v2.0.0 [31] was used to create neighbour-joining trees for each species based on whole genomes. Genomes were sketched using mash with default parameters prior to using mashtree. Mash distance was calculated for all isolates relative to the first genome entry for the group, in order to remove outliers with a distance > 0.1, which are likely to have had their species misclassified. Trees were subsequently rooted to outgroups and ultrametric trees were produced with the ape package in R [32] using the chronos function with smoothing parameters (λ) of 1 and 10. The “correlated”, “discrete” and “relaxed” models were used for each value of λ, which vary in the extent to which they permit adjacent parts of the tree to evolve at different rates. The tree with the highest log-likelihood for each species was used. Due to the large number of genomes and low prevalence of CRISPR-Cas, we did not perform this analysis for *S. aureus*.

The MCMCglmm package [33] was used to run Bayesian GLMs incorporating trees. The default weakly informative priors were used, and all models were run for 30,000 iterations with a burn in period of 20,000 iterations and a thinning interval of 5. Poisson GLMs were run for each species, with count of ABR genes as the response variable and CRISPR-Cas type or spacer count (in which case genomes with no CRISPR-Cas were removed) as a fixed effect. In cases where more than one CRISPR-Cas type was present, CRISPR-Cas type × total spacers was fit as an interaction. The small sample size for *S. epidermidis* with CRISPR-Cas systems (n = 54), meant we chose not to model the association between spacer count and ABR genes for this species. Prediction data frames for spacer counts were created up to the maximum count of spacers detected for that species, with the exception of *S. pyogenes*, for which the top value was an outlier at 33 so the next highest value of 17 was taken.

### Code availability and reproducibility

The snakemake pipeline used to calculate genome lengths and detect CRISPR-Cas systems, ABR genes, *intI1*, ICEs and plasmid replicons, as well as R scripts used for producing trees and statistical analysis for this work are available at https://github.com/elliekpursey/crispr-pathogens. Snakemake v5.18.1 [34] and Python v3.8.3 with Biopython v1.78 [35] were used for this work.

## Results

### Distribution of CRISPR-Cas systems, ABR and mobile elements across species

Genome analyses of ~40,000 genomes of *Pseudomonas aeruginosa*, *Acinetobacter baumannii*, *Neisseria meningitidis*, *Staphylococcus aureus*, *Staphylococcus epidermidis*, *Streptococcus pyogenes*, *Francisella tularensis*, *Mycobacterium tuberculosis*, *Neisseria gonorrhoeae*, *Enterococcus faecium* and *Enterococcus faecalis* revealed a large variety of CRISPR-Cas system types, ABR genes, ICEs and plasmids (Supp. Fig. 1). CRISPR-Cas systems were identified in every species except *N. gonorrhoeae*, which was therefore excluded from subsequent analyses.

Overall, type I-C, I-E, I-F, II-A, II-B, II-C, III-A, IV and V-A systems were represented across the dataset and ABR genes were detected in every species. However, *F. tularensis* was positive only for its native β-lactamase (FTU-1) [36], *M. tuberculosis* isolates were almost only ever positive for its intrinsic *blaA*, *aac*(*2*’)-*Ic* and *erm*(*37*) genes, and *N. meningitidis* had a maximum of 1 ABR gene, so these species were not considered in downstream analyses. The proportion of genomes with CRISPR-Cas varied across species from around 0.75 for *M. tuberculosis* to around 0.005 for *S. aureus* (Supp. Fig. 2). The number of repeats per genome also varied considerably between system type and species (Supp. Fig. 3), reaching as high as ~200 in I-F systems in *A. baumannii*, and ~150 in III-A systems of *M. tuberculosis*. However, for all other types this number was typically below 50.

Plasmids were detected in >80% of genomes for species in the *Staphylococcus* and *Enterococcus* genera, and were also prevalent in *S. pyogenes* (~25%), *P. aeruginosa* (10%), and *A. baumannii* (~3%) (Supp. Fig.1). As genomes were assembled to varying levels within the dataset, it is assumed some plasmids will have been missed using this method. ICEs were also common in these species, except for *A. baumannii*, ranging from 93% of *E. faecalis* genomes having at least one ICE to 15% of *E. faecium* genomes. Finally, *intI1* was detected in ~20% of *P. aeruginosa* and *A. baumannii* genomes.

### No clear association between genome size and CRISPR-Cas presence/absence

Consistent with previous work, *P. aeruginosa* genomes with CRISPR-Cas were on average smaller than those without the system. This was in fact the largest difference we saw, with predicted lengths of 6,579,416 ± 2885 bp and 6,692,831 ± 2601 bp for genomes with and without CRISPR-Cas, respectively (Supp. Fig. 4). However, the opposite was true in other species such as *A. baumannii* and *S. aureus*. For example, an *A. baumannii* genome with CRISPR-Cas was predicted to be 4,028,659 ± 5523 bp long, whilst one without had a predicted length of 3,976,319 ± 2181 bp, a difference of ~50,000 bp overall. Typically, differences between CRISPR-Cas positive and negative genomes were < 55,000 bp.

### CRISPR-Cas presence is associated with fewer acquired ABR genes and mobile elements, but more ICEs

We modelled the association between CRISPR-Cas presence or absence and the counts of ABR genes, as well as their vectors. Predictions are presented for elements which are found in each species in Figure 1 and all model estimates are presented in Supplementary Table 4. Across species, every additional ABR gene led to a 0.08 reduction in the probability of having a CRISPR-Cas system (*p* = 9.2E-18). The direction and strength of this trend varied by species, with positive associations between probability of CRISPR-Cas presence and ABR gene count in *S. epidermidis* and *P. aeruginosa* (Fig. 1A). CRISPR-Cas presence appeared to have little association with plasmid replicon count across species (Fig. 1B), whilst for *intI1*, a negative association was present in *P. aeruginosa* but not *A. baumannii*. For genomes possessing ICEs, positive associations were detected between ICE count and ABR gene count in *P. aeruginosa* and *S. pyogenes*.

**Figure 1 –.**
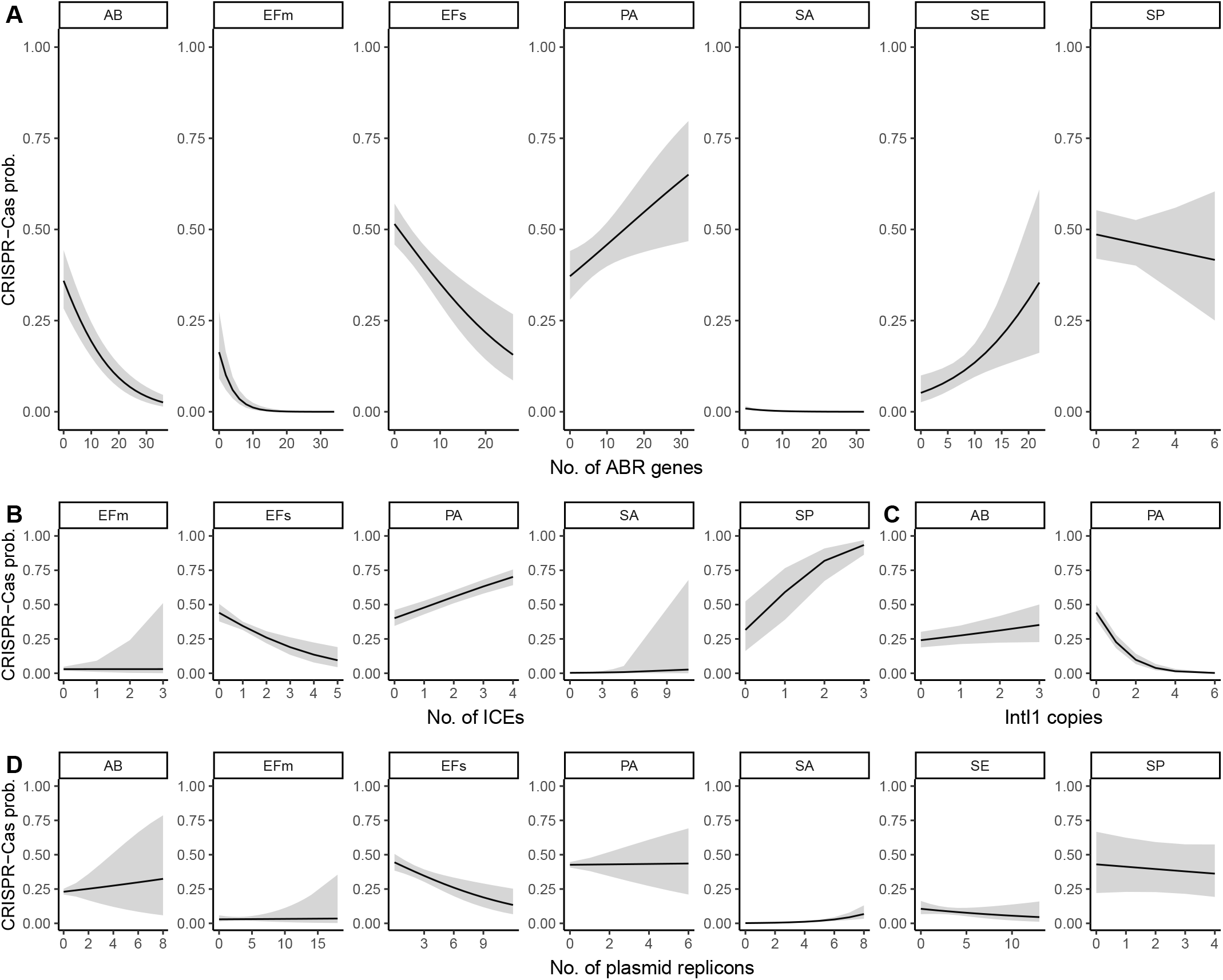
Predicted probability of CRISPR-Cas presence (y-axis) from binomial GLMs for A) ABR gene count and numbers of the potential ABR vectors B) plasmids, C) *intI1* (integrons) and D) ICEs. Predictions are presented for elements which are found in each species. Error bars show 95% confidence intervals.

### CRISPR-Cas presence is associated with fewer ABR genes across system types in individual species, controlling for genetic distance

To further explore the link between ABR genes and CRISPR-Cas, we made within-species models controlling for genetic distance, to look at the effect of CRISPR-Cas type and spacer count (Fig. 2). The overall trends detected are largely similar to those presented for across species comparisons in Fig. 1, in that there is generally a negative association between the presence of a CRISPR-Cas system and ABR counts. Across species, lacking a CRISPR-Cas system was associated with a higher predicted count of ABR genes (Fig. 2a). In some cases the size of this effect is striking; for example, an *E. faecium* genome lacking CRISPR-Cas is predicted to have 12 ABR genes compared to only 5 ABR genes for one possessing a type II-C system. However, in most cases this effect is more modest, with a reduction of 1 or 2 ABR genes when CRISPR-Cas is present. There are, however, two notable exceptions to this trend. Firstly, *Streptococcus pyogenes* type II-A systems were associated with a small but nonsignificant increase (0.44, 95% CI = 0.35 - 0.56) in the number of predicted ABR genes relative to genomes without CRISPR-Cas (0.33, 95% CI = 0.28 - 0.37). Secondly, both type I-C and IV systems are consistently associated with an increased count of ABR genes.

**Figure 2 –.**
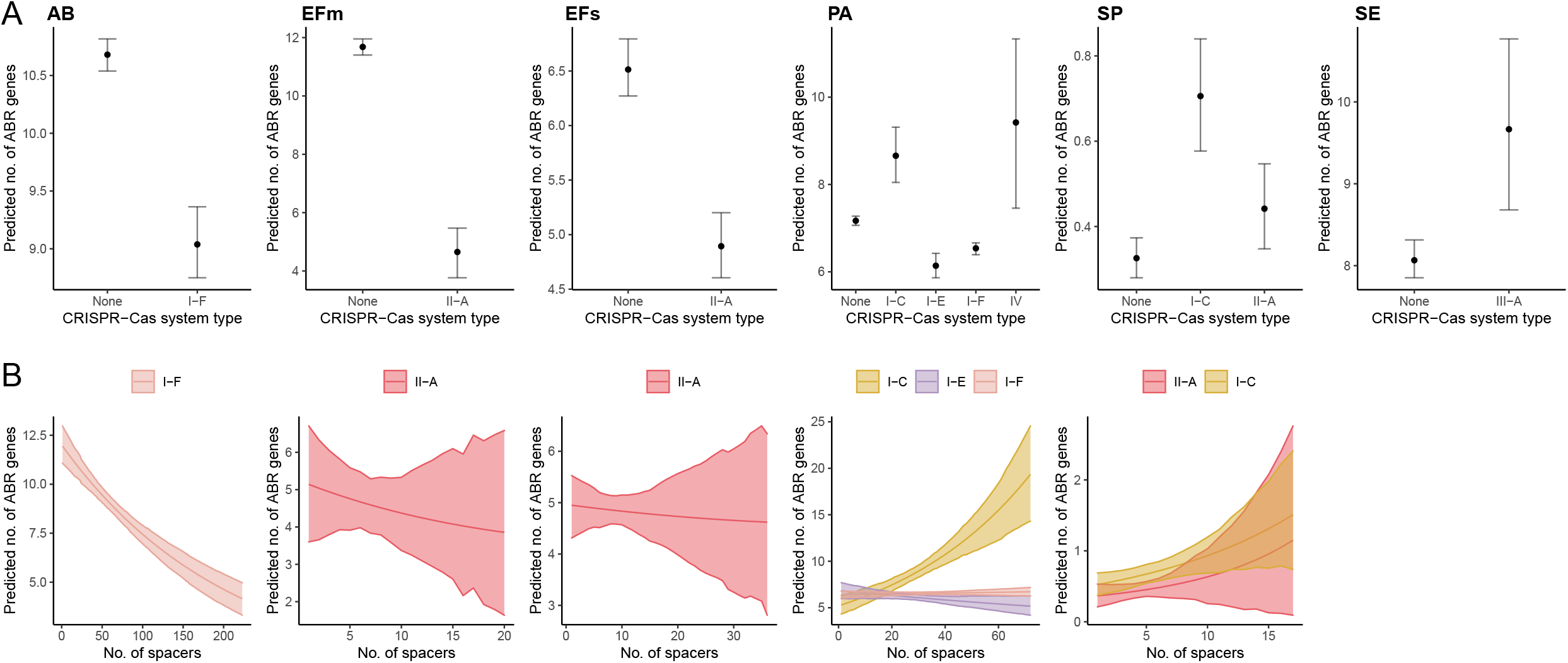
Prediction plots from Bayesian Poisson GLMs of ABR gene counts for A) CRISPR-Cas type and B) count of spacers. Each column represents an individual species. Error bars show 95% credible intervals.

Next, we modelled the number of ABR genes in response to the count of spacers (for those genomes that possess CRISPR-Cas; Fig. 2b). Here we did not find strong associations for most CRISPR-Cas system types. Exceptions were the type I-F system of *A. baumannii*, which showed a dramatic reduction in ABR gene count with increasing spacer count, and the type I-C system of *P. aeruginosa*, which interestingly showed the opposite trend. We did not model this association for type IV spacers due to the limited number of genomes (n = 20) in this category.

To expand on this dataset, we looked at the targets of all spacers by searching them against virus, ICE and plasmid databases (Supp. Fig. 5). As expected, the majority of the ~285,000 spacers detected (90%) did not have a match to any of these elements. Overall, the majority of known spacer targets are viruses (n = 24,402). However, we also detect many spacers targeting ICEs (n = 3,002) and plasmids (n = 1,416).

### Presence of spacers targeting ABR-carrying MGEs is negatively associated with ABR genes

As we detected many spacers matching potential vectors of ABR (n = 1,948), we wanted to determine whether there was an association between the presence of these spacers and an absence of ABR genes. We modelled the number of ABR genes in response to the number of spacers targeting MGEs that were positive or negative for ABR genes. The number of predicted ABR genes per genome was consistently lower when spacers targeted MGEs carrying ABR than when they targeted those that did not carry ABR, a trend that was universal across species (Fig. 3).

**Figure 3 –.**
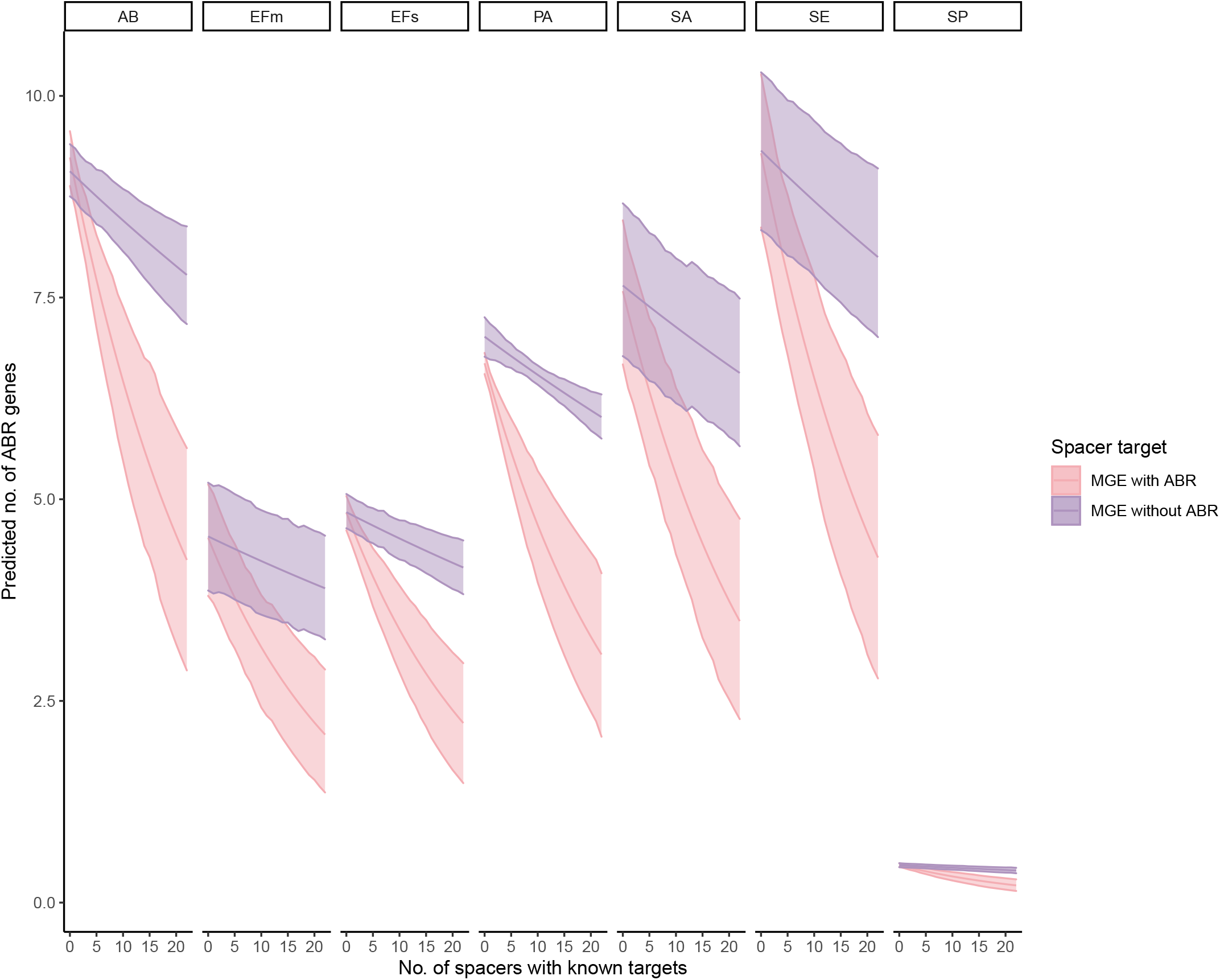
Prediction plot overlaying two Bayesian Poisson GLMs of ABR gene counts according to counts of spacers (with known targets) targeting MGEs with and without ABR, faceted by species. Error bars show 95% credible intervals.

## Discussion

We found little evidence for genome length as a marker of CRISPR-Cas blocking HGT in this collection of bacterial pathogens. Genome reduction is a process that is common in pathogens relative to their less-pathogenic relatives [37]. Bacteria may streamline their genomes in order to save energetic resources and remove genes that have become deleterious, or reductions in genome length may happen as a result of many other evolutionary processes that occur during the transition to pathogenicity. For example, in *P. aeruginosa*, strains adapted to the cystic fibrosis lung are frequently auxotrophic, as they are able to utilise amino acids produced by the host [38]. The extent of mobile DNA in genomes also reduces during the transition from facultative to obligate lifestyles, potentially due to reduced opportunities for their spread in more intracellular niches [39]. Therefore, this measure may be detecting potential adaptation to a pathogenic lifestyle or particular ecological environments, as well as a concurrent change in HGT frequencies.

However, we found compelling evidence that CRISPR-Cas can block the transfer of MGEs. We focused on the spread of ABR and its potential vectors due to their global importance, and identified a negative association between the count of ABR genes and the probability a genome will have a CRISPR-Cas system, which was repeatable across most CRISPR-Cas system types and species. Further supporting this hypothesis, we found genomes with spacers targeting vectors of ABR did indeed have fewer ABR genes. We did not detect strong trends for plasmid replicons, potentially because 1) genomes were assembled to varying levels and sequenced using different technologies within our dataset and many plasmids will have been missed and 2) the database we used was built based on plasmids found in *Enterobacteriaceae*, and has been shown to fail to predict plasmids in many species [40].

When looking at specific CRISPR-Cas types within species, we found that ABR genes were lower in frequency if CRISPR-Cas was present in most cases. Two exceptions were type I-C and IV systems, which were associated with more ABR genes. These system types may be driving the positive association we see between CRISPR-Cas presence and ABR genes in our multi-species model for *P. aeruginosa*. This result is also interesting in light of the fact that both of these system types are present on MGEs. Type I-C systems occur on conjugative elements in *P. aeruginosa* and have been found to correlate positively with the presence of particular ABR genes [11], whilst type IV CRISPR-Cas systems are usually found on plasmids, where they can mediate inter-plasmid competition [41]. CRISPR-Cas system types that are more mobile may co-occur with ABR genes found on mobile elements more frequently. Interestingly, the species for which we detected a positive association between ICEs and CRISPR-Cas presence in our multi-species model (*P. aeruginosa* and *S. pyogenes*) were also the only species to possess type I-C systems. We also saw positive associations between the count of ABR genes and the number of type I-C spacers in *P. aeruginosa*, suggesting that this system may even allow the genome to accumulate more ABR genes, and could have a role in competition between MGEs rather than host genome defence. However, much remains to be learned about these widespread but relatively understudied type I-C systems to fully explain this trend [42].

We do not see strong negative associations between the total count of spacers and the presence of ABR genes across most species and CRISPR-Cas system types. As the majority of spacer targets that we could identify were viruses, which are rarely vectors of ABR [43], the large number of anti-phage spacers and comparative rarity of spacers targeting ABR vectors may obscure any trend we would otherwise see here. Alternatively, the lack of clear association may be explained by the tendency for arrays to include more inactive spacers as they increase in size, as the most recently acquired spacers are most active [44]. Therefore, genomes with more spacers and larger arrays may include more spacers with no activity in removing potential vectors of ABR. An exception to this trend was *A. baumannii*, which had a strong reduction in ABR gene count with increasing spacer count in its type I-F system. Our spacer target analysis found that, unusually, more spacers in this species targeted plasmids than viruses; therefore, there could be some specialisation towards blocking plasmid acquisition in this system, a trend which has also been suggested for the type I-F system in *E. coli* [45].

We looked for spacers that targeted vectors of ABR and found they were fairly common. An increase in the count of spacers targeting vectors of ABR had a striking effect on reducing the predicted count of ABR genes. Particularly compared with the weaker association between the count of spacers targeting vectors without ABR and ABR genes, this evidence strongly suggests CRISPR-Cas is an important barrier to vectors of ABR, thereby blocking the acquisition of ABR genes themselves. Many ICEs, phages and plasmids remain to be characterised, and once our databases of these elements expand, this hypothesis will become more easily testable using genomic data.

This study supports the hypothesis that CRISPR-Cas system loss occurs where ABR acquisition is beneficial. In this case, the presence of CRISPR-Cas may be selected against in populations that are exposed to antibiotics, where the survival benefits of acquiring ABR genes outweigh the cost of phage predation or MGE maintenance. This has important implications for novel treatments, as multidrug-resistant bacteria may represent an immunocompromised population. In recent years, increasing attention has been given to antibiotic alternatives such as phage therapy [46,47] and CRISPR-Cas antimicrobials delivered using phage-like or plasmid elements [48,49]. Our results support the use of these interventions in cases where antibiotics do not work, as they are likely to be most effective in multidrug-resistant bacteria.

Our results also raise the question of why such trends have not been detected in previous work asking similar questions, particularly when looking across species. The strength of our work is primarily the use of more powerful and appropriate statistical tests, as well as controls for genetic distance. We selected the model with the best fit to the data, accounting and controlling for the effects of multiple predictors under one modelling framework as well as testing interactions between predictors. For example, we could test how the probability of a genome having a CRISPR-Cas system varies based on the number of ABR genes it has, whilst controlling for the number of ICEs, integrons and plasmid replicons in the genome overall. We could also control for interactions; for example, this allowed us to detect both positive and negative relationships between CRISPR-Cas presence and genome length for different species within the same model, avoiding some of the pitfalls of multiple testing. Such approaches are routinely used in evolution and ecology studies to account for the variability of datasets [50], and while suitable for these types of genomic comparisons, they are less commonly used in this context. Importantly, we also have access to a much larger dataset of genomes and improved computational tools since the publication of some previous work, which allows us greater statistical power to uncover associations between CRISPR-Cas and ABR genes.

## Conclusion

We found that looking both across and within-species, CRISPR-Cas system presence was associated with a reduction in counts of ABR genes. In addition, where spacers targeted MGEs that carry ABR, a concurrent reduction in ABR genes was seen. These results have promising implications for the delivery of novel technologies to combat ABR such as phage therapy and CRISPR-Cas antimicrobials, which rely on bypassing bacterial immune systems to kill cells. In fact, they may be ideal for this purpose, if the most multidrug-resistant strains are also the most immunocompromised.

## Acknowledgements

We would like to thank MD Sharma for his invaluable help with accessing computational resources needed for this work.

## Funding

EP is supported by a PhD studentship equally funded by the European Research Council (ERC-STG-2016-714478 – EVOIMMECH) and the College of Life and Environmental Sciences, University of Exeter. SVH acknowledges support from the Biotechnology and Biological Sciences Research Council (BB/S017674/1 and BB/R010781/10). FLP acknowledges support from Utrecht Exposome Hub of Utrecht Life Sciences (www.uu.nl/exposome), funded by the Executive Board of Utrecht University.

## Competing Interests

We have no competing interests.

## Notes

https://github.com/elliekpursey/crispr-pathogens

### Competing Interest Statement

The authors have declared no competing interest.

