## Supplementary figures and images for "CRISPR-Cas is associated with fewer antibiotic resistance genes in bacterial pathogens"

### Supplemental Figure 1

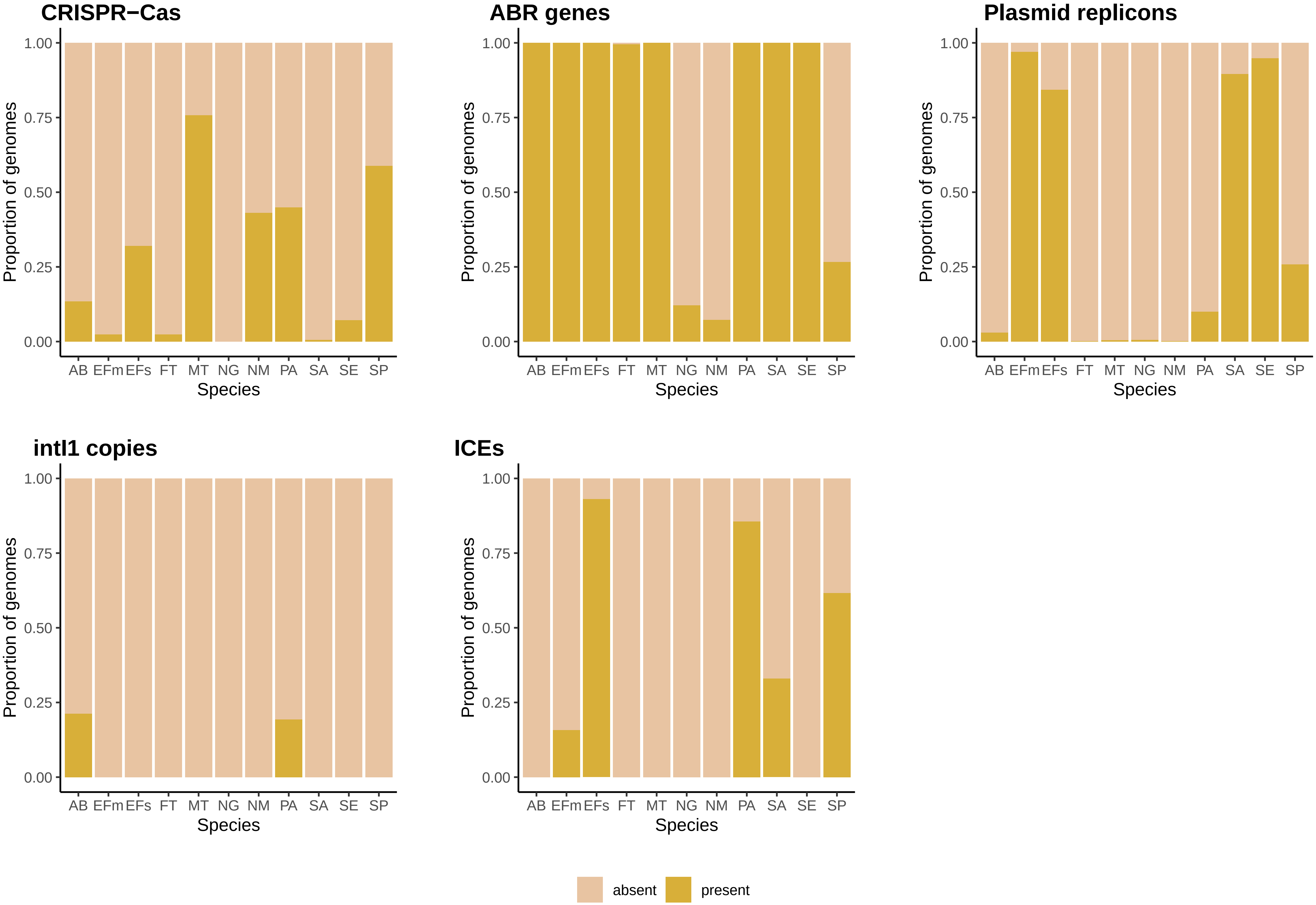

### Supplemental Figure 2

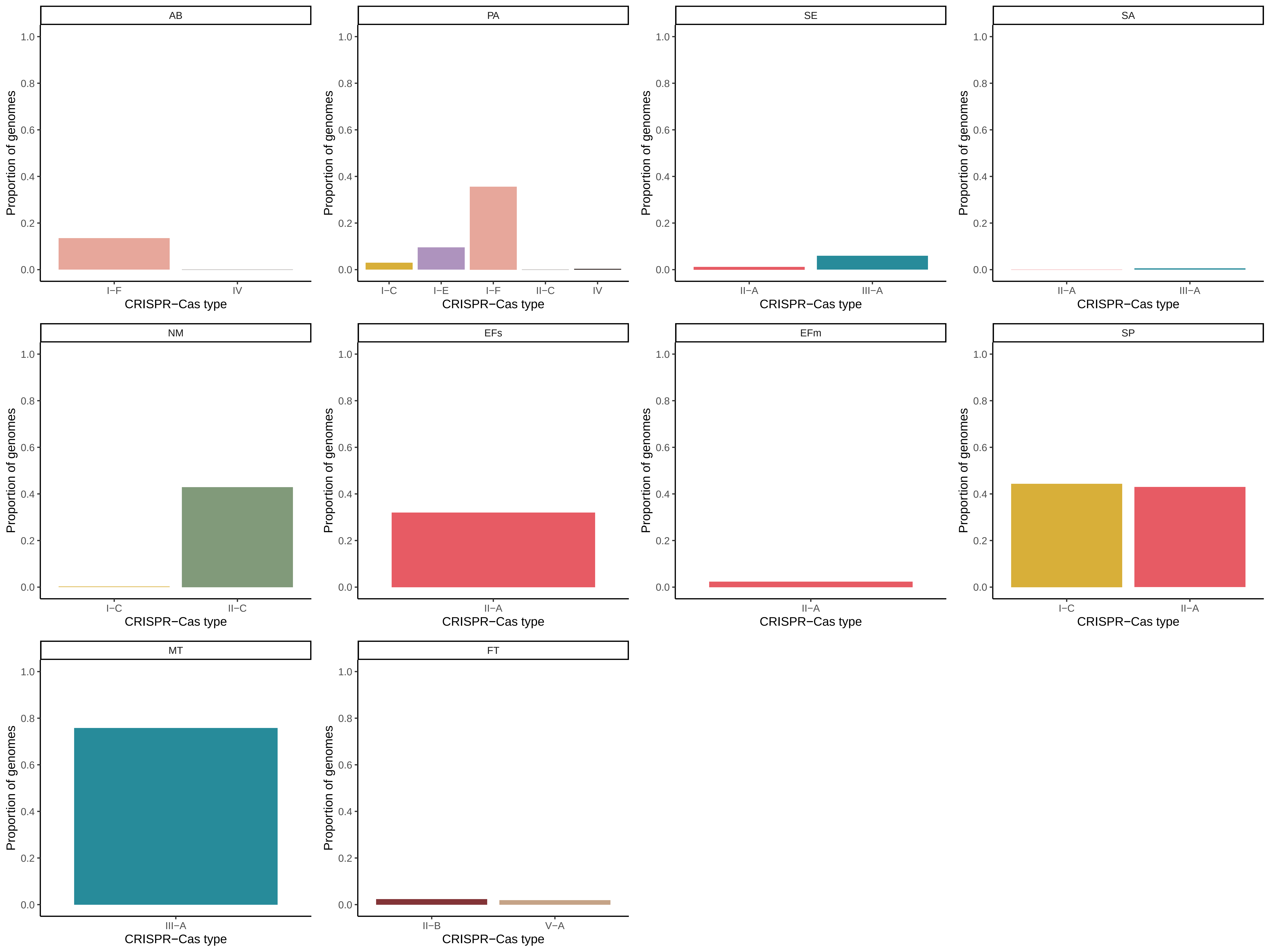

### Supplemental Figure 3

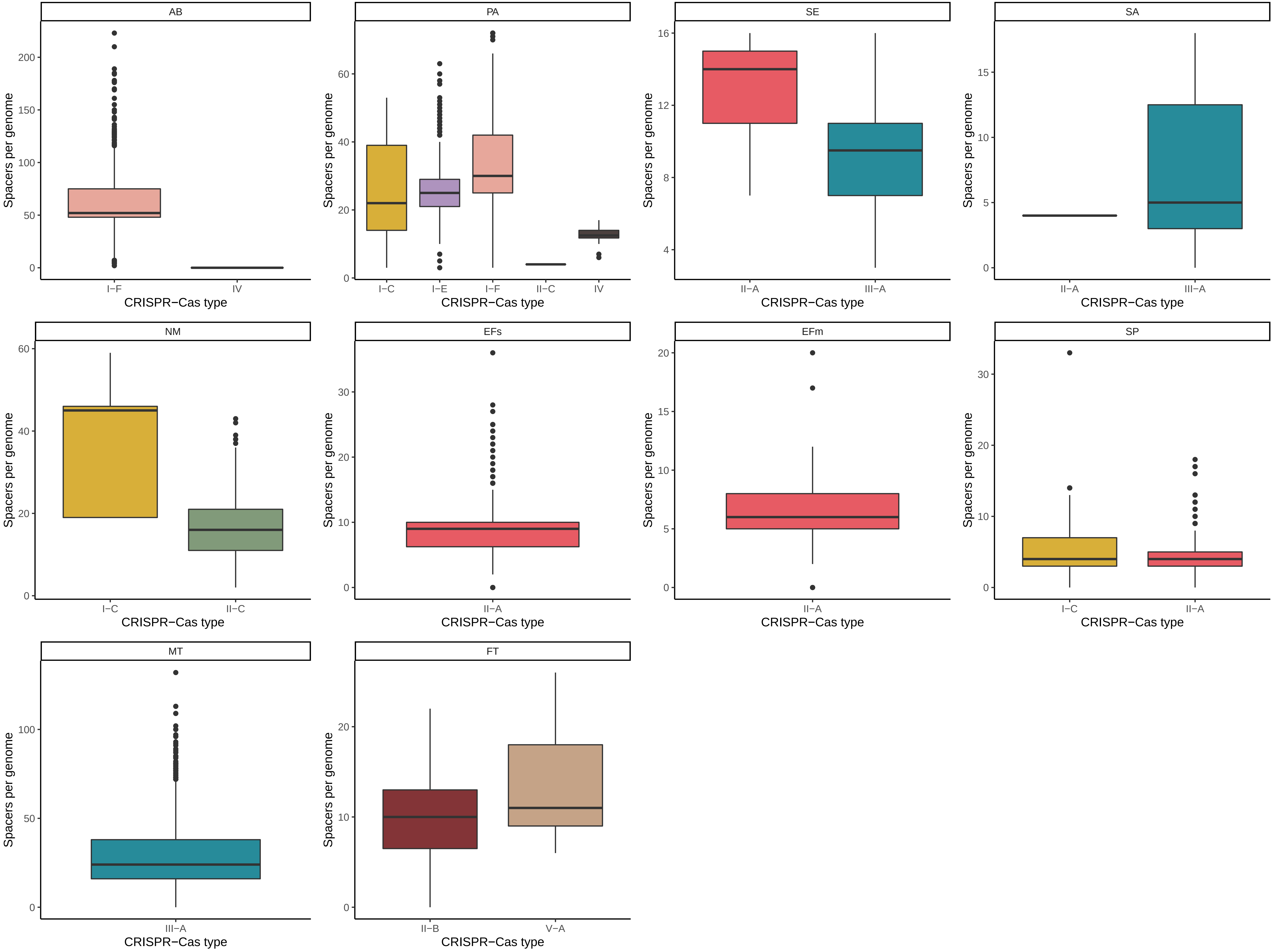

### Supplemental Figure 4

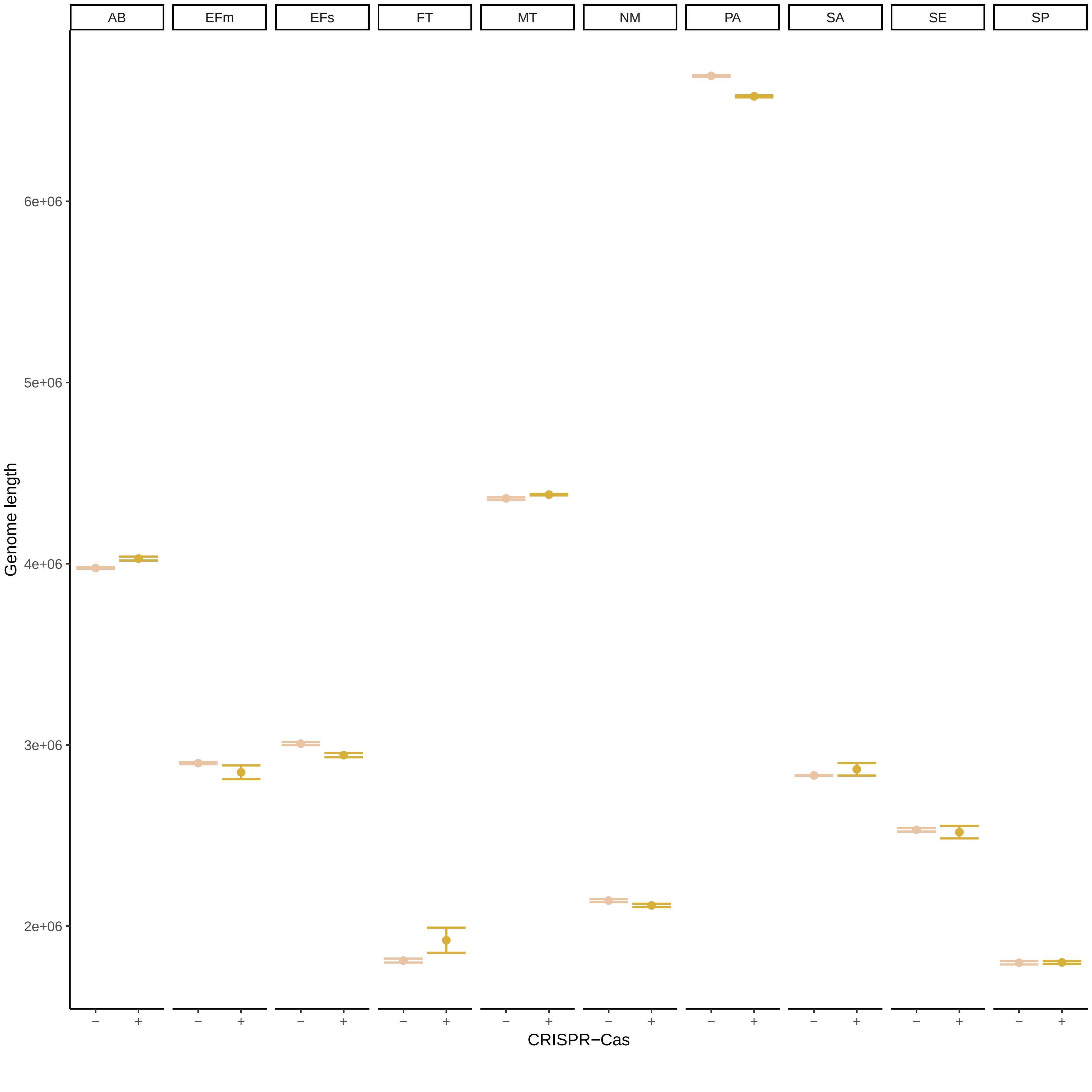

### Supplemental Figure 5

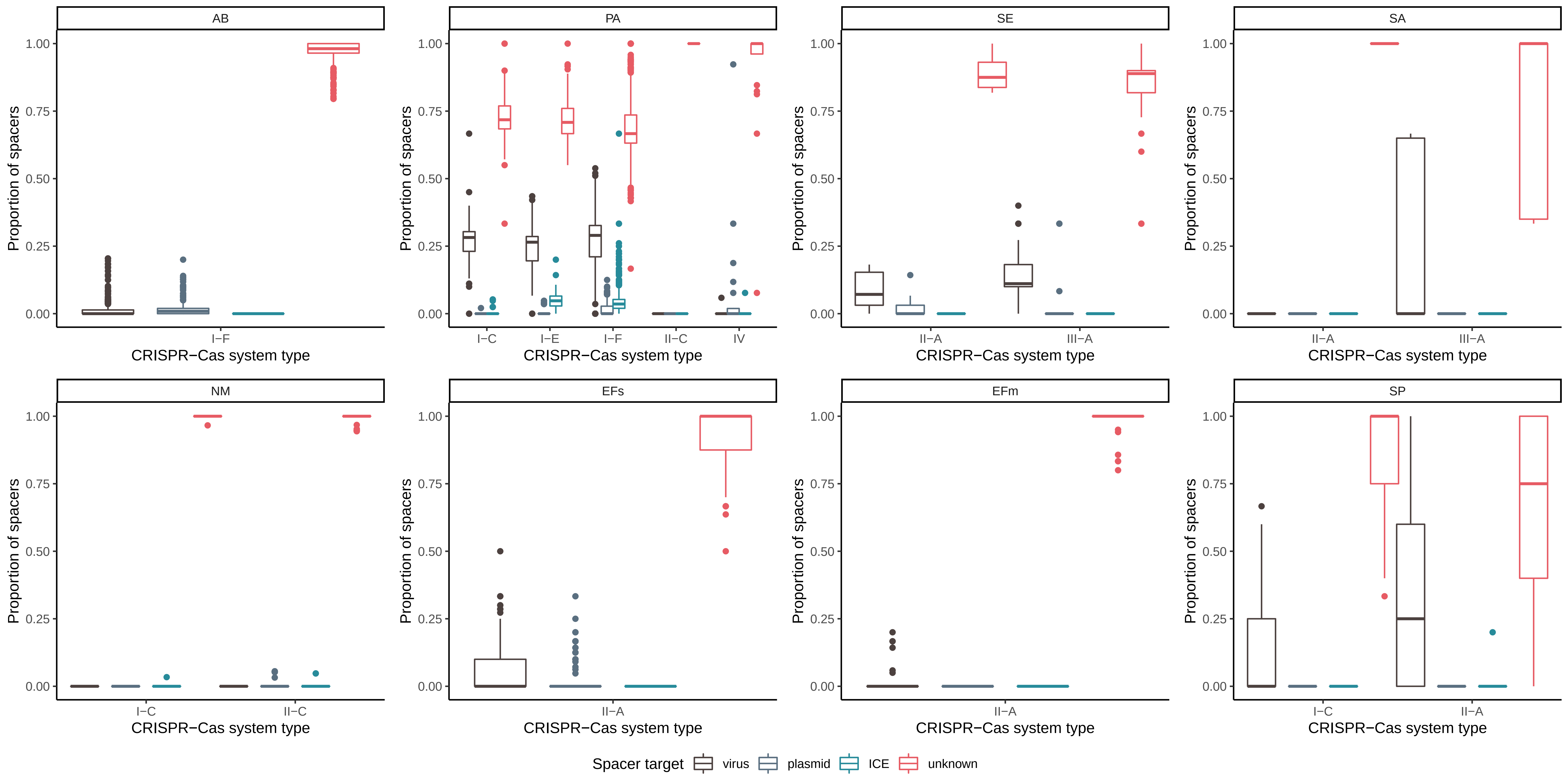
