## Supplementary figure and table legends for "CRISPR-Cas is associated with fewer antibiotic resistance genes in bacterial pathogens"

**Supplementary Material**

**Table legends**

Table S1 – AIC selection tables for linear model of genome lengths with fixed effects of CRISPR-Cas presence/absence (binomial), species (categorical) and an interaction term between the two. The fixed effect presence/absence from the model is indicated by +/- in the appropriate column. The prediction plot from the selected model (row 1) is presented in Supplementary Fig. 4.

Table S2 – AIC selection tables for binomial generalised linear model of CRISPR-Cas presence/absence with fixed effects of number of ABR genes, ICEs, *IntI1* copies and plasmid replicons, as well as species as fixed effects. The fixed effect presence/absence from the model is indicated by +/- in the appropriate column. The prediction plot from the selected model (row 1) is presented in Fig. 1.

Table S3 – Model estimates for linear model of genome lengths with fixed effects of CRISPR-Cas presence/absence (binomial), species (categorical) and an interaction term between the two. *Predictions were not made for this species, as it lacks CRISPR-Cas.

Table S4 – Model estimates for binomial generalised linear model of CRISPR-Cas presence/absence with fixed effects of number of ABR genes, ICEs, *IntI1* copies and plasmid replicons as well as species, and interaction terms between species and each other fixed effect.

Table S5 – Posterior distributions for individual species-specific Bayesian Poisson generalised linear models of ABR gene count with the fixed effect of CRISPR-Cas type (Prediction), and controlling for genetic distance. Intercept is always “None” i.e. no CRISPR-Cas system.

Table S6 - Posterior distributions for individual species-specific Bayesian Poisson generalised linear models of ABR gene count with fixed effects of CRISPR-Cas type (Prediction) and spacer count, and controlling for genetic distance.

Table S7 - Posterior distributions for Bayesian Poisson generalised linear model of ABR gene count with fixed effects of CRISPR-Cas type (Prediction) and proportion of spacers targeting MGEs that do not target ABR.

Table S8 – Posterior distributions for Bayesian Poisson generalised linear model of ABR gene count with fixed effects of CRISPR-Cas type (Prediction) and proportion of spacers targeting MGEs that target ABR.

**Figure legends**

Figure S1 – Proportion of genomes in each species with 1 or more CRISPR-Cas systems, ABR genes, plasmid replicons, intI1 copies and ICEs. AB, *Acinetobacter baumannii*; EFm, *Enterococcus faecium*; EFs, *Enterococcus faecalis*; FT, *Francisella tularensis*; MT, *Mycobacterium tuberculosis*; NG, *Neisseria gonorrhoeae*; NM, *Neisseria meningitidis*; PA, *Pseudomonas aeruginosa*; SA, *Staphylococcus aureus*; SE, *Staphylococcus epidermidis*; SP, *Streptococcus pyogenes*.

Figure S2 – Proportion of genomes possessing each type of CRISPR-Cas system, faceted by species. AB, *Acinetobacter baumannii*; EFm, *Enterococcus faecium*; EFs, *Enterococcus faecalis*; FT, *Francisella tularensis*; MT, *Mycobacterium tuberculosis*; NG, *Neisseria gonorrhoeae*; NM, *Neisseria meningitidis*; PA, *Pseudomonas aeruginosa*; SA, *Staphylococcus aureus*; SE, *Staphylococcus epidermidis*; SP, *Streptococcus pyogenes*.

Figure S3 –Number of repeats per genome for each CRISPR-Cas system type, faceted by species. AB, *Acinetobacter baumannii*; EFm, *Enterococcus faecium*; EFs, *Enterococcus faecalis*; FT, *Francisella tularensis*; MT, *Mycobacterium tuberculosis*; NG, *Neisseria gonorrhoeae*; NM, *Neisseria meningitidis*; PA, *Pseudomonas aeruginosa*; SA,

*Staphylococcus aureus*; SE, *Staphylococcus epidermidis*; SP, *Streptococcus pyogenes*.

Figure S4 – Prediction plot showing predicted genome length for CRISPR-Cas negative (-) and positive (+) genomes, faceted by species. Error bars show 95% confidence intervals. AB, *Acinetobacter baumannii*; EFm, *Enterococcus faecium*; EFs, *Enterococcus faecalis*; FT, *Francisella tularensis*; MT, *Mycobacterium tuberculosis*; NG, *Neisseria gonorrhoeae*; NM, *Neisseria meningitidis*; PA, *Pseudomonas aeruginosa*; SA, *Staphylococcus aureus*; SE, *Staphylococcus epidermidis*; SP, *Streptococcus pyogenes*.

Figure S5 – Proportions of spacers targeting viruses, plasmids, ICEs and with unknown targets, separated by CRISPR-Cas system type and faceted by species. AB, *Acinetobacter baumannii*; EFm, *Enterococcus faecium*; EFs, *Enterococcus faecalis*; FT, *Francisella tularensis*; MT, *Mycobacterium tuberculosis*; NG, *Neisseria gonorrhoeae*; NM, *Neisseria meningitidis*; PA, *Pseudomonas aeruginosa*; SA, *Staphylococcus aureus*; SE, *Staphylococcus epidermidis*; SP, *Streptococcus pyogenes*.
